# An in vitro model to study the naïve human peripheral blood mononuclear cell response to *Cryptococcus neoformans* infection

**DOI:** 10.1101/2020.04.08.032268

**Authors:** Katherine Pline, Simon A. Johnston

**Author notes:** Corresponding author: Simon Johnston.

## Abstract

*Cryptococcus neoformans* is a facultative intracellular pathogenic yeast which causes cryptococcal meningitis. Infection most commonly occurs via the lungs in humans and is cleared without clinical signs in the immunocompetent, but may cause life-threatening infection in immunocompromised. *C. neoformans* can be phagocytosed by host phagocytes, but may manipulate the intracellular niche of phagocytes for replication and dissemination. The interaction of macrophages with cryptococcal cells has been studied in detail but little is known about the interaction between human peripheral blood mononuclear cells (PBMC) and *C. neoformans*. PBMCs are rapidly recruited to the site of initial infection in the lung, peripheral tissues, and also respond to cryptococci that disseminate via the bloodstream. Therefore, deciphering the interactions between PBMCs and cryptococci is an important but neglected aspect of our understanding of the immune response during cryptococcal infection. Here, using time lapse imaging of primary human PBMCs *in vitro*, we are able to measure the PBMC response to cryptococci. Using this approach we find that naïve, undifferentiated human monocytes phagocytose cryptococcal cells, and that aggregates (swarms) of monocytes and T cells often form in response to engulfment of cryptococci. Interestingly, we find a correlation between the size of the PBMC aggregates and proliferative ability of the cryptococci within. While these aggregates slow intracellular cryptococcal growth, cryptococci are able to replicate within this niche and escape from PBMCs to replicate extracellularly. These results provide evidence for PBMC control of cryptococcal infection and provide a model for the *in vitro* study of cryptococcal granuloma biology.

## Introduction

*Cryptococcus neoformans* is a facultative intracellular pathogen that causes the fungal infection cryptococcosis (1, 2). It is estimated that globally over 223,000 people suffer from cryptococcal meningitis each year, resulting in 181,000 deaths (3). While this pathogen may infect people who have competent immune systems (4), *C. neoformans* mainly infects people who are severely immunocompromised, particularly people with late stage HIV infection/AIDS. Serological evidence shows that children without symptomatic cryptococcosis have antibodies which are reactive with cryptococcal proteins, suggesting control of this infection by the host immune system in the immunocompetent (5). Furthermore, there is some evidence for *Cryptococcus* being able to cause a latent infection with the reactivation of dormant cryptococci after extended periods of sub-clinical infection (6, 7). As with the majority of opportunistic infections normal immunity to cryptococcosis is not well understood, but limited clinical studies suggest a role for granulomas in the control of acute cryptococcal infection (8). Thus, *C. neoformans* may be effectively cleared by the host, lie dormant, or produce life threatening infection in the form of cryptococcal meningitis.

Interactions between *C. neoformans* and the host immune response are complex and appear to be largely governed by interactions between the pathogen and host phagocytes (9, 10). It is believed that *Cryptococcus* is inhaled as a spore or (desiccated) yeast from the environment and can be deposited in the alveoli of the lung where alveolar macrophages attempt to clear the infection (11). While macrophages have been shown to be important for the control of early infection (12, 13), *C. neoformans* may also exploit the phagocytic niche to replicate and disseminate infection. Indeed, murine infection with macrophages which contain intracellular cryptococci results in enhanced dissemination compared to infection with free fungal cells (14), illustrating a role for macrophages in cryptococcal dissemination. Furthermore, vomocytosis of fungal cells is associated with dissemination in vivo (15), and recent work suggests that *C. neoformans* likely colonises monocytes and macrophages to escape from the lung environment (16). Despite these insights into the role of macrophages in cryptococcal pathogenesis, there is still much to learn about the immune response to cryptococcal infection. An immune population that has been largely neglected in the study of cryptococcosis are the peripheral blood mononuclear cells (PBMCs). Composed of both innate and adaptive immune cells, PBMCs are a unique population that serve both a patrolling and a response role to pathogenic threats throughout the body. The PBMC population contains cells which phagocytose foreign threats as well as cells which support these innate functions to provide robust control. Given their role in the immune response, it is likely that PBMCs are important in the control of cryptococcal infection at the initial site of infection, as well as in potential control of fungemia following dissemination from the lung and pathogenesis at sites of dissemination.

Several studies provide data concerning the potential PBMC response to cryptococcal infection, but varying cell incubation periods and ambiguity in the developmental stage of the assayed cells fail to capture the early response of naïve human PBMCs immediately after isolation. For example, in the presence of pooled human serum, Levitz and colleagues demonstrated PBMCs respond to cryptococcal infection by releasing TNF-α, and that monocytes isolated from this population are able to bind fungal cells more potently after six days of infection (17); however, this study failed to disclose the PBMC incubation period, and monocytes incubated for six days are likely to be at least partially differentiated into macrophages, thus failing to capture the immediate PBMC response. A later study investigated the long-term PBMC response to cryptococcal infection. Siddiqui and colleagues ‘pre-stimulated’ human PBMCs with heat-killed *C. neoformans* for twelve days before infecting cells with live *C. neoformans* for up to ten days (18). This study showed that pre-stimulation of human PBMCs induced better control over secondary live infection compared to unstimulated PBMCs in a manner that is dependent on fungal capsule size and IL-6 production (18). However, this long term of pre-stimulation followed by long periods of live infection fails to capture the immediate naïve PBMC response to cryptococcal challenge that is likely to occur after initial cryptococcal infection. While these studies have illustrated how peripheral immune cells interact with the invading fungal cells, the delayed introduction of the pathogen until after a period of immune cell differentiation means that the monocytes were likely at least partially differentiated into macrophage-like cells from the monocytic precursors. Furthermore, assays which utilise pathogenic pre-stimulation before live infection fail to address the response of naïve, freshly isolated PBMCs which are likely to encounter the pathogen during initial infection and dissemination in the periphery. Therefore, the initial PBMC response to cryptococcal infection remains largely unknown.

Here, by performing time lapse imaging on PBMCs infected with *C. neoformans* at <24 hours post-isolation, we show that cryptococci are phagocytosed by PBMCs and are able to replicate within the host cells *in vitro*. We demonstrate how we can measure phagocytosis, fungal cell growth, and vomocytosis within a naïve PBMC population. Using our approach, we show that PBMCs swarm in response to fungal intracellular replication, and that fungal growth is limited within the PBMC niche. These results show that an immunocompetent *in vitro* model of cryptococcosis results in granuloma-like aggregate formation that limits cryptococcal infection by reducing fungal growth, and provide a new opportunity for the field to understand the immune response to *C. neoformans* infection.

## RESULTS

### The vast majority of donor PBMCs co-incubated with cryptococci were CD14+ monocytes and CD3+ T cells

While PBMCs are often characterised by live by flow cytometry, we needed a non-invasive live imaging method to understand the makeup of our isolated PBMC population in culture. To do this we used directly fluorescent dye-conjugated antibodies to discern the immune cells composing the human blood samples at both 0 and 24 hours post infection (hpi). Antibody staining revealed that the majority of the cells present were CD14+ monocytes and CD3+ T cells (Fig. 1A). Additionally, CD3+ T cells exhibited a more uniformly circular appearance where CD14+ monocytes were larger and more heterogeneous, allowing them to be distinguished. At 0 hpi, both T cells and monocytes were largely adherent to the plate (Fig. 1A); however, at 24 hpi T cells were mostly non-adherent (Bennett and Breit, 1994). As expected, we did not observe neutrophils which are generally excluded by PBMC isolation techniques, but did observe a small number of platelets that are commonly co-isolated with PBMCs. Spontaneous aggregates of PBMCs were commonly observed throughout our assays. Aggregates appeared to be composed mainly of CD14+ monocytes. T cells contributed to aggregates at 0 hpi, and to a lesser extent at 24 hpi (Fig. 1A and 1B). There were a very small number of unlabelled cells, most likely natural killer or B cells. These results showed that our donor PBMCs were, as expected, almost entirely composed of CD14+ monocytes and CD3+ T cells.

**FIG 1.**
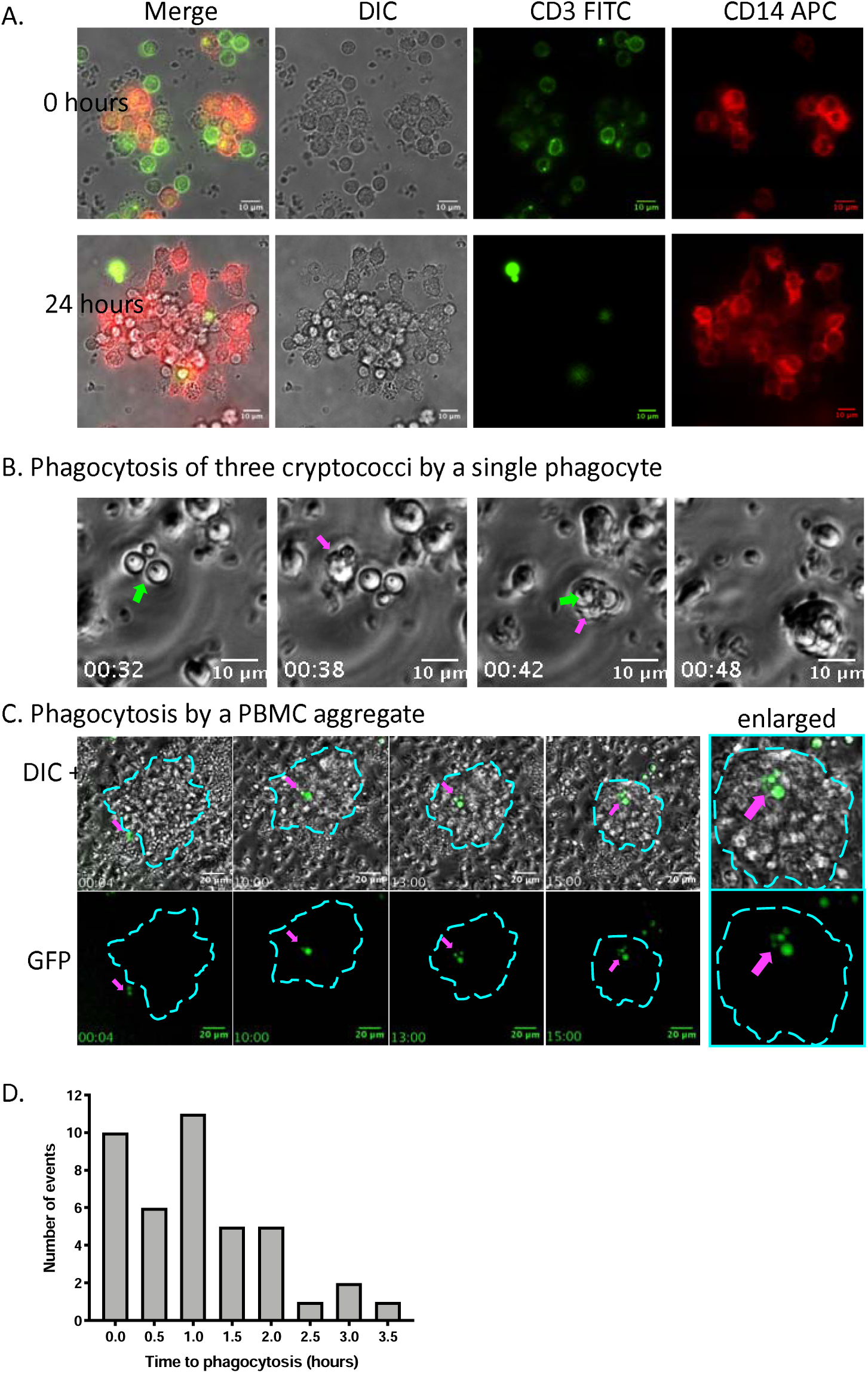
Immune cell subsets and uptake of fungal cells. (A) Human PBMCs were isolated for 24 hours and then antibody staining was performed on live cells to identify CD3+ T cells and CD14+ monocytes. Green FITC-labelled cells are T cells and red APC-labelled cells are monocytes. Imaging was performed at x60 magnification after 0 and 24 hours. Fewer T cells were adherent at 24 hours, though both subsets composed aggregates. (B) Phagocytosis of KN99 GFP *C. neoformans* (green arrow) by human peripheral monocyte (magenta arrow). Time is shown as hours:minutes. (C) Phagocytosis of *C. neoformans* by an aggregate of human PBMCs. Two cryptococci (shown in GFP and marked by magenta arrow) were phagocytosed by a pre-existing swarm of immune cells (outlined in cyan). As the experiment progressed the cryptococcal cells divided from 2 to 5 cryptococcal cells. TIme is shown as hours:minutes. (C) The time each fungal cell was phagocytosed was determined and plotted as a histogram. Bars represent the number of cryptococci phagocytosed at each time point. Three repeats were performed, each representing a different PBMC donor. 41 starting cryptococci were followed.

### CD14+ peripheral monocytes, but not CD3+ T cells, phagocytosed cryptococcal cells

To understand how cryptococci interact with PBMCs, time lapse imaging was performed to follow individual cryptococci and observe their interactions with PBMCs. At approximately 24 hours after PBMC isolation, cells were infected with opsonised KN99 GFP *C. neoformans*. Cryptococci were phagocytosed by individual monocytes (Fig. 1B) and by monocytes within PBMC aggregates (Fig. 1C), though not all cryptococci were phagocytosed. Additionally, cryptococci were phagocytosed as individual cells or as small groups of up to four cryptococci (Fig. 1B). Most phagocytosis events occurred immediately within two hours of infection (Fig. 1D) and additional phagocytosis of cryptococci occurred rarely throughout the duration of the assays.

### PBMCs form swarm-like aggregates in response to cryptococci

Time lapse analysis of cryptococcal infection showed the formation of PBMC aggregates as described previously (18). Frequently, after phagocytosis additional PBMCs migrated to the phagocyte which possessed intracellular cryptococci, reminiscent of swarming behaviour. These aggregates were variable; many were only small and transient, while other aggregates persisted and formed much larger masses (Fig. 2A). While we found that PBMCs were able to swarm to form a mass of cells around phagocytosed cryptococci, we did observe cryptococci being phagocytosed by cells in pre-existing masses (Fig. 2A – right panel *iv*). Aggregates containing cryptococci sometimes fused with additional aggregates that did not contain fungal cells (Fig. 2B).Intriguingly, larger aggregates that met usually did so only transiently before splitting apart, each retaining their ‘original’ intracellular cryptococci with any resultant daughter cells. Not all swarms persisted throughout the course of the 24 hours, and individual phagocytes (including those containing cryptococci) were seen to split away from masses at apparently random times throughout.

**FIG 2.**
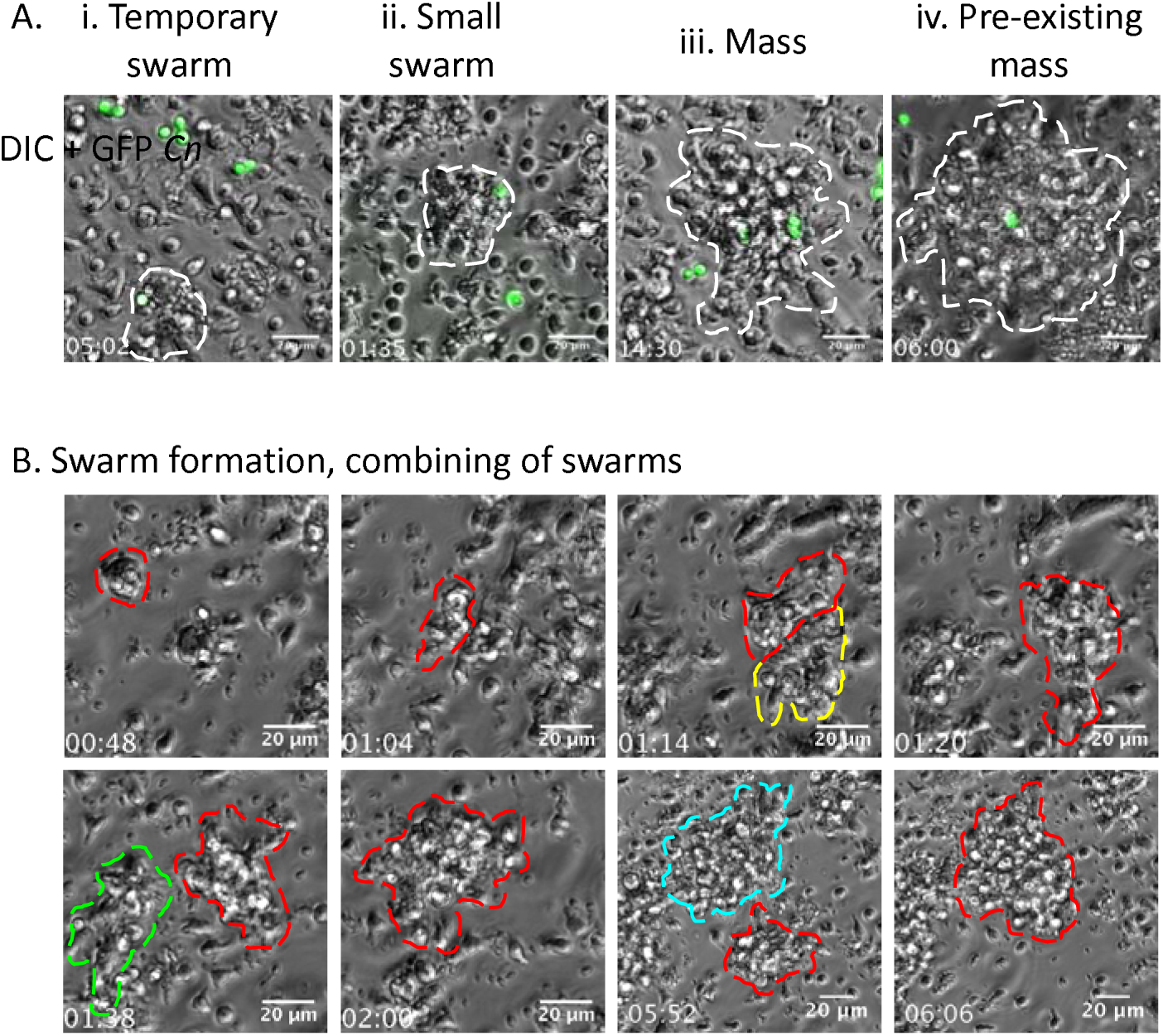
PBMCs form aggregates in response to cryptococcal infection. Human PBMCs were infected with KN99 GFP *C. neoformans* one day after isolation. Time lapse imaging was performed at x20 magnification for up to 24 hours. (A) Different types of PBMC aggregates were seen during infection. Cryptococci are shown in green and aggregate boundaries are marked by white dashed lines. (i) Small temporary swarm of PBMCs around a phagocyte containing intracellular cryptococci. Swarm formed only briefly then dissolved to leave a single monocyte containing a fungal cell. (ii) Swarm which persisted around an infected phagocyte. (iii) Formation of a larger mass with intracellular cryptococci. (iv) Phagocytosis of a cryptococcal cell by a pre-existing mass. Cryptococcal cell was phagocytosed by the swarm of by a phagocyte in the margin, as opposed to first being phagocytosed by a monocyte before the formation of a swarm. (B) The formation of a swarm around an infected phagocyte in time. Swarm boundaries are marked with coloured dashed lines, and red represents the development of the swarm around the original phagocyte. The phagocyte was met with a small number of PBMCs to form a small swarm (second frame), which then combined with subsequent swarms (yellow, green, cyan lines) to produce a large swarm. While cryptococci are within they are not represented here. Time is shown as hours:minutes.

### PBMC aggregate size was not related to the number of intracellular fungi but increased in proportion to fungal growth

Characteristics of the PBMC aggregates were quantified in order to understand the role of these PBMC aggregates during cryptococcal infection. Most aggregates formed within four hours of phagocytosis (Fig. 3A). Out of 40 aggregates followed, 11 (27.5%) showed no fungal replication, whereas 29 (72.5%) harboured replicating intracellular cryptococci. Because the presence of pathogens may induce immune activation, the size of the aggregates was measured and compared to the number of cryptococci within. We hypothesised that if there were immune activation in response to phagocytosis of the pathogen, that more intracellular cryptococci would result in the formation of a larger swarm size. A relationship between aggregate size and the number of cryptococci could also suggest that cryptococci recruit more PBMCs to exploit as a niche. To test this, the size of each swarm was measured each time a cryptococcal cell was replicated. The size of the aggregates was compared to the number of intracellular cryptococci contained within each, and showed that there was no clear relationship between the swarm size and the number of cryptococcal cells within (Fig. 3B). Indeed, while many aggregates increased in area, some decreased in size. Though there was no clear relationship between the size of aggregates and the number of cryptococcal cells they contained at any given moment, there was a positive linear relationship between the relative fold change of the intracellular cryptococci and the maximum size of the swarm (Fig. 3C; linear regression p = 0.0014, R^2^ = 0.283). Here, the maximum size of the swarm increased as the cryptococci within displayed enhanced proliferative ability. This suggested two intriguing possibilities: that either larger aggregates were more permissive to intracellular proliferation, or that monocytes in which cryptococci were replicating were stimulating PBMC swarming. This may indicate that cryptococci which were more prolific intentionally recruited a higher number of PBMCs for use as an immune cell niche for replication. These results may alternatively suggest that increased swarming of PBMCs in response to prolific cryptococci was due to immune recruitment in order to control the infection. For comparison, the size of aggregates lacking intracellular cryptococci was also measured at the start and end of the time lapse assays and revealed no population trend for increase or decrease in size (Fig. 3D), supporting our finding that individual factors, such as cryptococcal growth, were most likely driving any difference rather than a simple single activation step to swarming. In addition, the inability of all ‘empty’ swarms to increase in time supports the idea that the increase in PBMC mass size with increasing fungal fold change may be related to the replication of cryptococci contained within, and may not be due merely to a general swarming behaviour.

**FIG 3.**
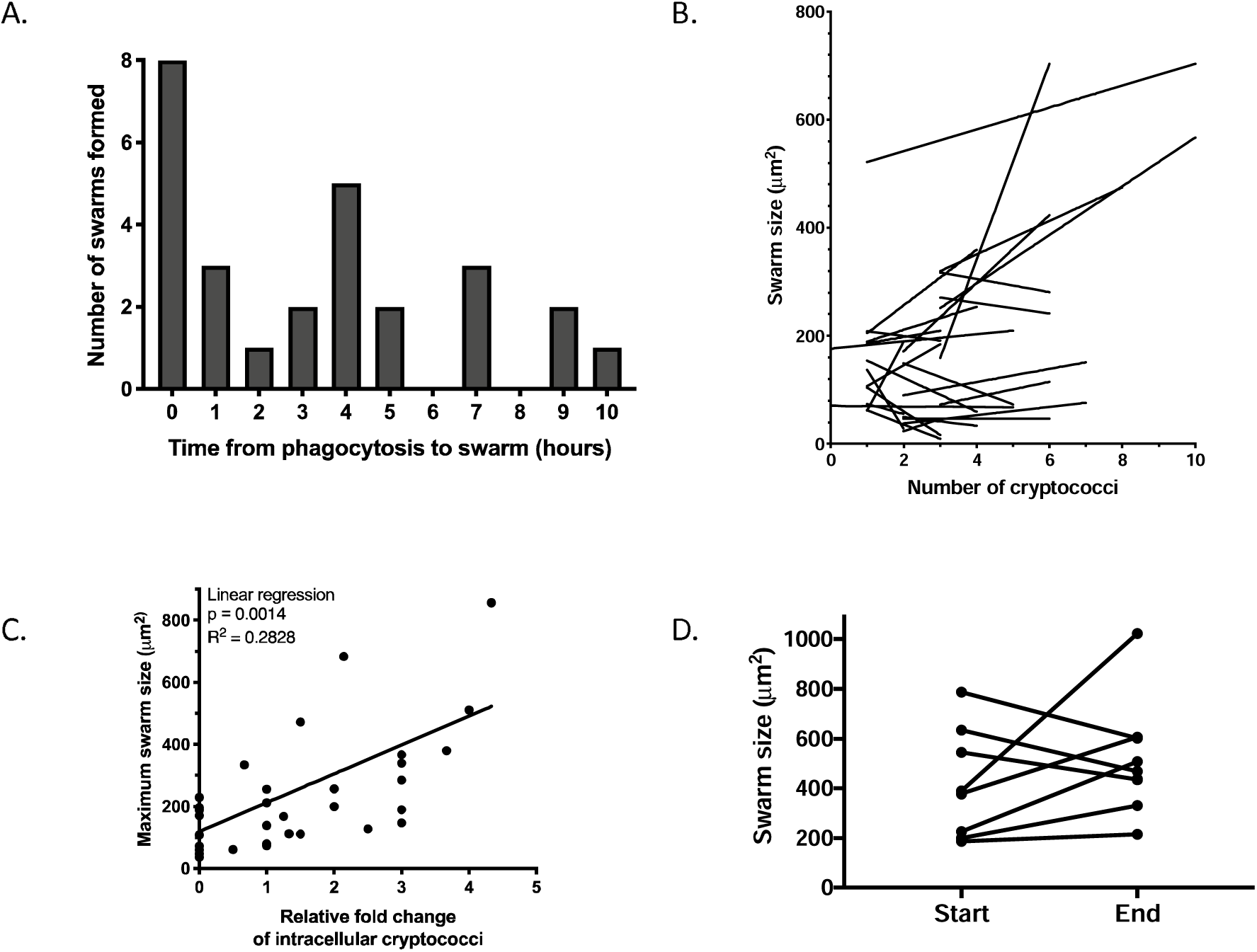
PBMC swarm behavior in response to cryptococcal infection. (A) Time between the phagocytosis of cryptococci and the formation of PBMC swarms. Swarms usually formed almost immediately in less than an hour. However, swarms could form throughout the experiment, though the majority aggregated within 4 hours of phagocytosis. (B) The area of the swarm in relation to the number of cryptococcal cells within. The size of the swarm was determined by drawing around the swarm and measuring the area in μm^2^. Measurements were made each time a division occurred. Each line is representative of one phagocytosis event leading to the formation of a swarm. (C) The relative fold change for each intracellular cryptococcal cell was measured and compared to the maximum size the swarm reached throughout the experiment. There was a positive linear relationship between the fungal relative fold change and the size of the mass (linear regression p = 0.0014, R^2^ = 0.2828). (D) Measurement of the size of swarms which did not contain cryptococci but were within the same well during cryptococcal infection. Swarm size was measured at the start and end of time lapse videos. Few of these types of aggregates existed and remained within the field of view. Those that did showed variable changes in size. Three repeats representing different donors were performed.

### Intracellular cryptococci showed reduced growth within human PBMCs

To distinguish a protective or facilitative role of PBMC swarming in controlling cryptococcal replication, individual fungal cells were followed over the duration of 24 hours to quantify the growth rates of intracellular (Fig. 4A), extracellular (Fig. 4B), and vomocytosed (Fig. 4C) fungi. While most cryptococci within phagocytes did replicate (Fig. 4A), those which were extracellular had a faster growth rate (Fig. 4B) with the doubling time shorter for cryptococci which were in the extracellular environment compared to those within phagocytes or swarms (Fig. 4D and 4E; one-way ANOVA p < 0.0001 when comparing intracellular doubling time to other conditions). In fact, cryptococci which were extracellular replicated twice as fast as those which were intracellular (mean doubling times 4.39 ± 1.6 hours for extracellular cryptococci, 9.34 ± 3.9 hours for intracellular cryptococci). Confirming the growth supressing intracellular environment, cryptococci that had been intracellular but vomocytosed from phagocytes or masses grew more quickly after escaping from PBMCs compared to intracellular cryptococci (Fig. 4D and 4E; mean doubling time 5.0 ± 3.5 hours) and supported our hypothesis that swarms were forming in response to intracellular growth. In addition, relative fungal fold change over time (growth rate) was also calculated to compare proliferative ability of fungi amongst conditions. We found that cryptococci which were intracellular show a reduced ability to replicate compared to those which were extracellular (Fig. 4E mean relative fold change 2.62 ± 1.77 for intracellular cryptococci, 11.58 ± 8.39 for extracellular cryptococci; 0.16 ± 0.11 replications per hour for intracellular cryptococci, 0.74 ± 0.42 for extracellular cryptococci). Vomocytosed cryptococci did not replicate as much as cryptococci which were extracellular from the start, suggesting that there may be some negative effect of vomocytosis on replication, or that transferring from an intracellular to an extracellular environment may require an adjustment phase before replication. However, this is unlikely because doubling times of vomocytosed and extracellular cryptococci were closely comparable. Finally, we found no difference between the doubling times of extracellular cryptococci and those in media alone without the presence of PBMCs (Fig. 4D and 4E; one-way ANOVA p = 0.0567). While this suggests that over the time considered here there was no extracellular killing or cytotoxic functions of PBMCs against cryptococci, the slightly higher doubling time of extracellular cryptococci in the presence of PBMCs suggests there may have been some slight inhibitory effect on growth.

**FIG 4.**
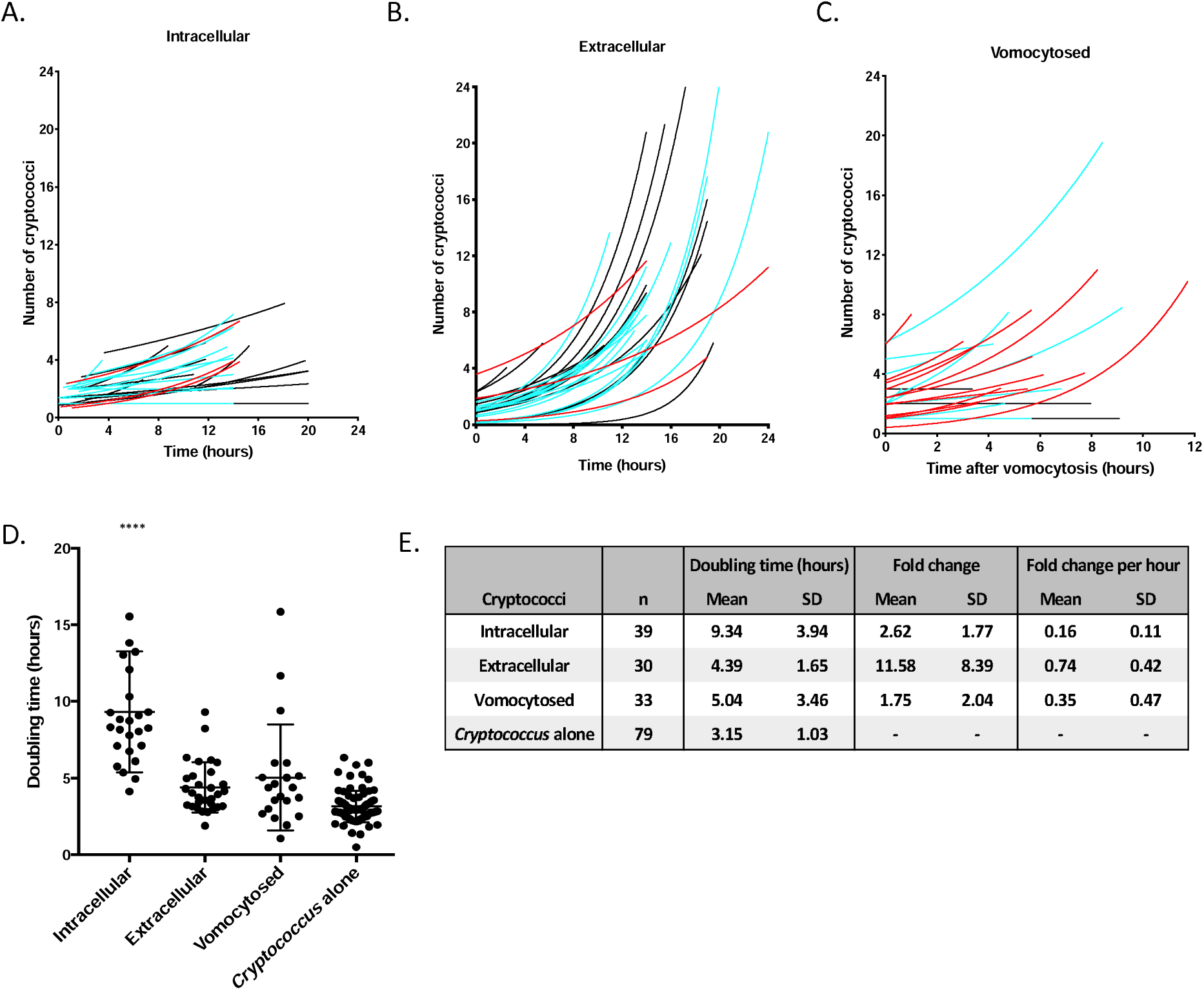
Replication of cryptococcal cells during PBMC infection. Three repeats were performed with different colours representing different donors. (A) The growth of individual cryptococci which were intracellular within phagocytes and PBMC swarms. (B) Quantification of the growth of individual extracellular cryptococci in the presence of human PBMCs. (C) The growth of individual cryptococci which had vomocytosed from phagocytes and PBMC masses. (D) The doubling times of cryptococci which were intracellular, extracellular, vomocytosed, or in SF RPMI media alone. Means and standard deviations are represented. The doubling time was calculated by fitting an exponential curve. Doubling time of intracellular cryptococci was significantly longer than other conditions (one-way ANOVA p < 0.0001 when comparing the mean doubling time of each condition to the doubling time of intracellular cryptococci). Cryptococci which were vomocytosed also had a longer doubling time than cryptococci which were alone in media in the absence of PBMCs, though the significance is left off the graph for the sake of clarity (one-way ANOVA p = 0.0048). (E) Descriptive statistics comparing the doubling times between cryptococci in different niches. N represents the number of cryptococci followed, with mother-daughter buds counted as one. Fold change of cryptococci is also shown, though for comparison the fold change normalised to the time followed is also given.

### Cryptococci can escape from PBMC aggregates and individual monocytes

Cryptococci are able to exit from host cells via vomocytosis (20, 21). Cryptococci escaped from monocytes via vomocytosis, and also escaped from aggregates either within an individual monocyte or via vomocytic expulsion (Fig. 5A). Out of 215 individual cryptococci followed, 100 (46.5%) vomocytosed during the 24-hour period. Of the remaining cells followed, 102 (47.4%) remained intracellular, while 13 (6%) disappeared due to death or were censored from the analysis due to an inability to confirm continuous analysis throughout the time lapse. Thus, while human PBMCs limit cryptococcal replication rate, they also fail to contain intracellular cryptococci as nearly half of all fungal cells escaped from the host intracellular niche. Additionally, some cryptococci appeared to produce large polysaccharide capsules (Fig. 5B) but we did not observe the formation of titan cells (22).

**FIG 5.**
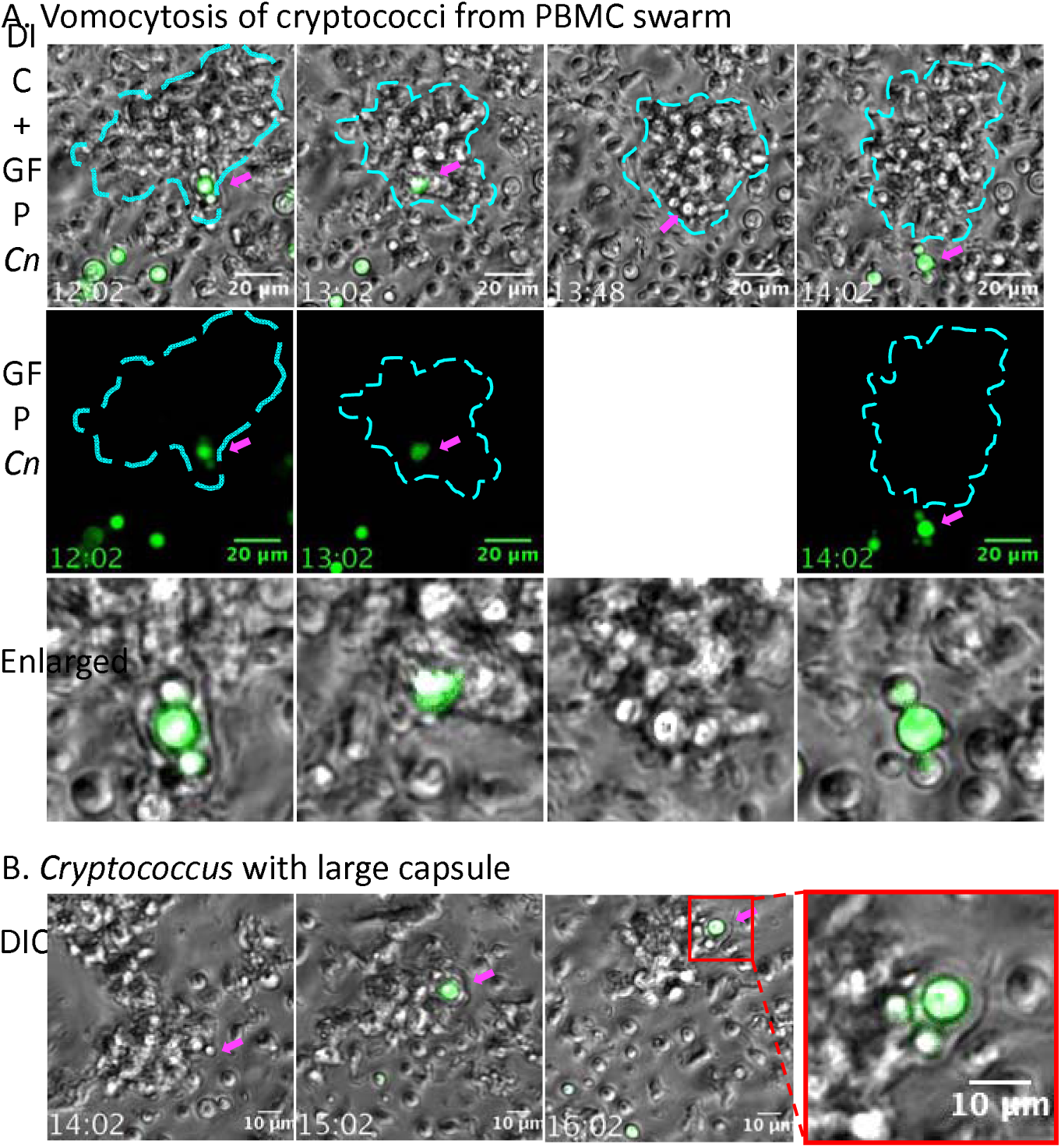
Cryptococci escaped from PBMC masses and produced large capsules. Human PBMCs were infected with KN99 GFP *C. neoformans* and followed for 24 hours via time lapse imaging. Behaviour of cryptococci within PBMCs was noted. (A) Cryptococci were able to escape both from individual phagocytes and from PBMC masses. They were vomocytosed as cryptococcal cells, or escaped the swarm in phagocytes. The vomocytosis of three joined cryptococcal cells is pictured, with cryptococci in green marked by the magenta arrow. The PBMC mass boundary is marked by cyan dashed lines. One panel is missing because GFP images were taken only once per hour, and this time point was between GFP image captures. Enlarged images are given below to illustrate intracellular and extracellular states (bottom panel). (B) The formation of an extended capsule formed around a GFP cryptococcal cell during its inhabitation of a PBMC swarm. The magenta arrow marks the location of the GFP cryptococcal cell. Throughout the time lapse this fungal cell struggled at the edge of the swarm yet failed to escape. Time is shown as hours:minutes.

## Discussion

Collaboration amongst immune cells to respond to pathogenic threat is important for control of infection, and understanding these immune responses provides insight into infection progression and eventual outcome. The macrophage immune response to cryptococcal infection is widely studied, but the interactions between cryptococci and PBMCs has been largely disregarded. PBMCs are a robust immune cell cohort that are likely to encounter cryptococci both at the site of infection in the lung at the onset of infection, as well as in the peripheral tissues during cryptococcal fungemia. As such, the PBMC response to *C. neoformans* is likely important for initial containment of infection, as well as control in the periphery before dissemination to the central nervous system. Here, we showed that after phagocytosis of cryptococci by monocytes, T cells and monocytes formed immune aggregates that slowed fungal growth compared to cryptococci in the extracellular environment. Size of aggregate formation was associated with intracellular growth of cryptococci within monocytes. However, these interactions failed to halt cryptococcal replication, and cryptococci were able to escape PBMC containment in this immunocompetent model of infection. These results show how peripheral monocytes and T cells respond to infection before monocytic differentiation into macrophages, and illustrate that this fungal pathogen is capable of immune exploitation and escape.

Phagocytosis of pathogens is a crucial component of the innate immune response. Time lapse analysis after PBMC infection with cryptococci showed that monocytes phagocytosed cryptococci almost immediately after encounter, usually within two hours. Rapid phagocytosis by monocytes illustrates their ability to engulf cryptococci before differentiation into macrophages; this is important as undifferentiated monocytes are likely to encounter cryptococci both within the lung environment as well as in the periphery during dissemination, proving an innate phagocytic response at crucial points of cryptococcal pathogenesis.

While phagocytosis of invading pathogens is imperative for control of infection, the behaviour of phagocytes after engulfing invading pathogens further determines the progression of infection. For example, recruitment of additional immune cells to contain the microbe is often necessary for immune control of infection. Here we showed that after phagocytosis of cryptococci by monocytes, additional PBMCs frequently swarmed to the infected monocyte. Events like this have been described previously where, after pre-stimulation of PBMCs with heat-killed cryptococci, secondary infection with live *C. neoformans* resulted in PBMCs swarming around engulfed cryptococci (18). While this previous study focused on the immune response after pre-stimulation of potentially differentiated monocytes, in the current study we showed that naïve, freshly-isolated PBMCs formed aggregates around infected monocytes, illustrated the PBMC response after the introduction of a primary Cryptococcus infection *in vitro*, and showed that swarming was related to intracellular growth of cryptococci in monocytes. These results suggest that at the initiation of cryptococcal infection in vivo, PBMCs may swarm to form aggregates around the invading fungal cells.

The formation of PBMC aggregates may serve to contain the cryptococcal infection. Many infections result in the production of granulomas for the containment of infection, and granulomas have been described for cryptococcal infection (7, 8, 23). Murine studies have shown that cryptococcal granulomas in the lungs were densely packed with mononuclear cells and macrophages with the addition of leukocytes, resulting in dense aggregates of organised immune cells (24, 25). In humans, cryptococcal granulomas are also characterised by densely organised monocytes, macrophages, and multinucleate giant cells containing intracellular cryptococci, as well as CD4+ lymphocytes (8, 23). Because the immune cell structures described in these studies are similar to the PBMC aggregates observed here, these PBMC aggregates are analogous to cryptococcal granulomas and present a tool for the study of granuloma-like aggregate formation *in vitro*.

The function of granulomas is to limit proliferation and dissemination, and to contain the pathogen if killing is hindered (26). However, as *C. neoformans* may manipulate the intracellular niche for replication and dissemination (1, 14, 15, 27–29), it is important to interpret the role of these aggregates with caution. While aggregates slowed cryptococcal proliferation compared to extracellular cryptococci, fungi still managed to proliferate within the intracellular niche. The relationship between fungal fold change and maximum swarm size suggested that PBMCs swarmed to highly prolific cryptococci to contain infection. This is in contrast to a phenomenon observed during *Mycobacterium marinum (Mm)* infection of zebrafish. During this bacterial infection, *M. marinum* induced granuloma formation for niche exploitation and bacterial intracellular replication (30), and recruitment of uninfected macrophages enabled serial macrophage invasion and propagation of infection (31). In our assays, because PBMC swarmed to control prolific infection, the major effect of PBMC swarm formation during *in vitro* cryptococcal infection appears to be control of cryptococcal growth. Despite this, the high number of cryptococci which managed to escape from monocytes and swarms suggests that these aggregates were unable to effectively contain cryptococci and prevent dissemination.

Though cryptococci were able to escape PBMC containment, monocytes were able to engulf the pathogen and slow replication, illustrating some inhibitory effect on cryptococcal growth. This is important as it illustrates that after introduction of cryptococcal infection naïve human PBMCs are able to engulf the *C. neoformans* and hinder replication, but that some cryptococci are able to escape the immune cells to replicate extracellularly. Additionally, for the first time this study illustrates and measures the swarming of naïve PBMCs during the early immune response to cryptococcal infection, which is important for the control of cryptococcal growth after initial infection and after dissemination to the periphery, both of which are important steps for cryptococcal pathogenesis. Taken together, the results described here show how the introduction of cryptococcal threat induces response by human peripheral immune cells which are likely to be important for controlling infection both in the lung and during dissemination in the periphery.

## Materials and methods

### Ethics statement

Informed written consent was obtained before whole blood was collected from healthy human donors. The South Sheffield Research Ethics Committee (REC) provided ethical approval for the collection of blood for the study of monocyte derived macrophages (REC reference 07/Q2305/7).

### Strains and Media

*Cryptococcus neoformans* var. grubii, strain KN99 GFP was used for all assays. For long term storage cryptococcal cells were stored in MicroBank vials at −80°C. Cells were rescued by streaking onto YPD agar plates (YPD 50 g/L Fisher, 2% agar Sigma-Aldrich) and incubated for 48 hours at 28°C. Plates were stored at 5°C until use. For use cryptococci were suspended in 2 ml of YPD broth and cultured overnight at 28°C, rotating horizontally at 20 RPM. 1 ml of KN99 GFP *C. neoformans* suspension was removed from the overnight preparation and was washed and pelleted using a centrifuge (3300g for 1 minute). Cells were washed using this method three times in PBS. Cells were diluted 1/20 and counted using a haemocytometer, and prepared at a concentration of 10^5^ cells per ml in serum-free Rosewell Park Memorial Institute (RPMI) medium. For infection cryptococci were opsonised using 0.1 μl of 18B7 (anti-capsule monoclonal antibody) per 100 μl of culture and were left to opsonise on a rotator for one hour.

### Cell isolation

PBMCs were isolated by the technicians at the Royal Hallamshire Hospital Medical School. After collection of blood samples, whole blood was added to Ficoll-Paque in a sterile centrifuge tube at a ratio of 1:2 (Ficoll:Blood). This was centrifuged at 1500 rpm for 23 minutes before the plasma layer was discarded. PBS was added to PBMCs, and this was centrifuged at 1000 rpm at 4°C for 13 minutes. Supernatant was removed, pellet was re-suspended in PBS, and was centrifuged again. Pellet was then resuspended in RPMI containing 10% fetal bovine serum (Sigma-Aldrich) and 1 ml was added per well in a 24 well plate, or 3 ml was added to individual glass-bottomed dishes used for antibody staining. PBMCs were prepared at of 2×10^6^ cells per well. Cells from three different donors were used.

### PBMC infection

Cryptococci were cultured, washed, diluted, and opsonised as above. PBMCs were infected the morning after isolation (less than 24 hours). Serum-containing RPMI was removed from cells, and cells were infected with 1ml of KN99 GFP *C. neoformans* in serum-free RPMI. Imaging occurred immediately after infection, or 24 hours after infection for some antibody assays.

### PBMC antibody staining

2 μl of FITC mouse anti-human CD3 (BD Pharmigen Biosciences), 2 μl mouse anti-human CD14 APC (Invitrogen, MHCD1405), and 20 μg of human IgG were added to glass dishes containing PBMCs. 100 μl of 10^4^ opsonised cryptococci was added. These dishes were either imaged immediately to represent the PBMCs at the onset of time lapse assays, or were incubated for 24 hours before imaging to represent the PBMCs at the end of 24-hour time lapse assays. Dishes were imaged at x60 magnification using differential interference contrast (DIC), DAPI, GFP, and Cy5 channels.

### Microscopy

Nikon eClipse TI microscope and FIJI (ImageJ) version 2.0.0 analysis software were used for imaging and analysis. x20 phase imaging was performed using perfect focus to capture time lapse videos. DIC images were captured every two minutes, and GFP images were captured once per hour. x60 imaging was performed for antibody labelling assays.

### Fungal replication measurement

Individual cryptococcal cells were followed by their GFP signal for up to 24 hours. Replication was quantified by measuring the time to the appearance of each budding daughter cell. The number of cryptococci was plotted against time for individual starting cryptococci, and doubling time was determined by fitting an exponential curve in GraphPad Prism (version 7.0). The relative fold change of cryptococci was found by counting the number of total cryptococcal cells which resulted from a starting (or several starting) fungal cells at 0 hours post infection. Relative fold change was calculated by the following: [(final number of fungal cells – starting number of cryptococcal cells) / starting number of cryptococcal cells]. This was normalised to the hours followed by dividing this number by the number of hours a cell or collection of cells was observed.

### Aggregate size measurement

PBMC aggregates were measured at the start and end of time lapse assays, or when a division of fungal cells within was observed. Measurement was performed using FIJI version 2.0.0 by drawing around the margin of an aggregate and finding the area in μm^2^.

### Statistics

GraphPad Prism version 7.0c was used for statistical analyses. Normality was determined based on adherence to a Gaussian distribution and normality tests were not performed. 95% confidence intervals were used to assess statistical significance. Ordinary one-way ANOVAs were performed to compare the means of conditions. Multiple comparisons were performed, using Tukey’s test to correct for multiple comparisons. Linear regression was also performed in GraphPad Prism.

## Acknowledgements

We would like to acknowledge the University of Sheffield for financial contributions under the Postgraduate Research Student Publication Scholarship Scheme to KP, SAJ was supported by Medical Research Council and Department for International Development Career Development Award Fellowship MR/J009156/1. SAJ was additionally supported by a Krebs Institute Fellowship and Medical Research Council Centre grant (G0700091). We would like to thank the Faculty of Medicine, Dentistry and Health and the Department for Infection, Immunity and Cardiovascular Disease for further financial support. We would also like to thank Jonathan Kilby and members of his team at the University of Sheffield Medical School for isolation of human cells and Helen Marriott for providing CD14 and CD3 antibodies.

